# Machine learning based lineage prediction from AMR phenotypes for Escherichia coli ST131 clade C surveillance across infection types

**DOI:** 10.1101/2025.09.22.677784

**Authors:** Theodor A. Ross, Anna K. Pöntinen, Einar Holsbø, Ørjan Samuelsen, Kristin Hegstad, Michael Kampffmeyer, Jukka Corander, Rebecca A. Gladstone

## Abstract

2.

Rising antimicrobial resistance (AMR) in *Escherichia coli* bloodstream infections (BSIs) in high-income settings has typically been dominated by one clone, the sequence type (ST) 131. More specifically, ST131 clade C (ST131-C) is associated with fluoroquinolone resistance and extended-spectrum β-lactamases (ESBLs). Even though urinary tract infections (UTIs) are a known common precursor to BSIs, there is currently limited knowledge on the longitudinal prevalence of ST131-C in UTIs and, therefore, the temporal link between the two infection types. Leveraging available genomic and antimicrobial susceptibility test (AST) data for ciprofloxacin, gentamicin, and ceftazidime in 2790 *E. coli* BSI isolates, we trained random forest and Extreme Gradient Boosting (XGBoost) classifiers to predict if an *E. coli* isolate belongs to ST131-C using only AST data. These models were used to predict the yearly prevalence of ST131-C in 22942 UTI and 24866 BSI isolates from Norway. The XGBoost classifier achieved a prediction F1-score of over 70% on a highly unbalanced dataset where only 4.3% of the genomic BSI isolates belonged to ST131-C. The predicted prevalence of ST131-C in UTIs exhibited a similar annual trend to that of BSIs, with a stable infection burden for eight years after its rapid expansion, confirming that the persistence of ST131-C in BSIs is largely driven by ST131-C UTIs. However, a higher prevalence of ST131-C in BSIs (∼7%) compared to UTIs (∼4%) suggests a subsequent enrichment of ST131-C. Our study highlights how existing epidemiological knowledge can be supplemented by utilising extensive data from AMR surveillance efforts without genomic markers.

**Impact statement:** This study proposes a potential analysis method that leverages AST data, which is already regularly collected for AMR surveillance purposes. Using such data to approximate the population-wide prevalence of MDR clones, such as ST131-C, could allow for larger-scale, retrospective studies of its prevalence in a population than genomic-based methods at a significantly lower cost. Such a method could supplement existing knowledge and epidemiology study practices. We use the proposed method to find relationships between the prevalence of the important MDR *E. coli* clone ST131-C in UTIs and BSIs in Norway. These results suggest that monitoring and reducing MDR in UTIs could reduce the burden of this invasive clone in hard-to-treat BSIs.

**Data summary:** All AST data, clonal information for isolates with genomic data, and code used in this study can be found in the following repository: https://github.com/theodorross/EColi-UTI-Predictions. Only the clone and published metadata information is shown for the UTI data shared by Handal, Kaspersen *et al*.

## 5. Introduction

Antimicrobial resistance (AMR) and multidrug resistance (MDR) in *Escherichia coli* bloodstream infections (BSIs) have been reported to be increasing in Norway and worldwide (Pöntinen et al. 2024; Gladstone et al. 2021; MacKinnon et al. 2021). Up to 67% of BSIs have been reported to have a urinary infection focus (Mehl et al. 2017; Abernethy et al. 2017; Day et al. 2016), making trends in urinary tract infections (UTIs) of importance for understanding the current BSI burden. However, limited data on the clonal composition of both UTIs and BSIs prevents the relationship between clonal prevalence in these two infection types from being explored (Day et al. 2016). The rise in AMR and MDR in BSIs is primarily driven by the expansion of a single clonal lineage, sequence type (ST)131. This clone has emerged as the single largest contributor of fluoroquinolone resistance and extended-spectrum β-lactamases (ESBLs) in *E. coli* BSIs in Norway during the last couple of decades (Gladstone et al. 2021). The clone is composed of four major sub-clades A, B, C1 and C2, which differ in their AMR profiles. ST131 C clades (ST131-C; C1 and C2) are uniformly fluoroquinolone-resistant and account for most (59%) ESBL-positive isolates in Norwegian BSIs. ST131 has also been shown to be an important cause of hard-to-treat UTIs (Wang et al. 2021; Li et al. 2023; Kudinha et al. 2013; C. M. Liu et al. 2018). Directly determining clonal prevalence over time usually requires molecular typing of isolates. This, in turn, necessitates that surveillance isolates are systematically stored, and significant resources are required for processing large longitudinal sample sets. Antibiotic susceptibility testing (AST) profiles, both the level of resistance and the exact combination of different antibiotic classes an isolate is resistant to, are strongly associated with clones, to the extent that the AMR profile of *E. coli* can be predicted based on the clone it belongs to (MacFadden et al. 2020). Therefore, we hypothesised that existing AST data could be a resource-efficient alternative for assessing the prevalence of important MDR clones.

Tandem advancements in whole genome sequencing (WGS) technologies and the field of machine learning (ML) have been leveraged in attempts to further the study of AMR in microbes. In addition to phenotypic testing for AMR, genomic sequence information can be used to predict resistance properties of bacterial isolates. One option is to use BLAST-based methods to compare a query genome to a curated database of AMR genes, such as CARD, AMRFinderPlus, and ResFinder (Alcock et al. 2023; Feldgarden et al. 2021; Florensa et al. 2022). These tools offer easily explainable categorical predictions but would fail if previously unknown genetic loci confer a resistance phenotype or contribute to a higher minimum inhibitory concentration (MIC).

ML is a powerful tool for analysing structure in datasets and has proven itself better equipped for detecting previously unknown resistance genotypes with classical techniques, such as tree-based and linear prediction models, applied to WGS data from multiple bacterial species (Deelder et al. 2019; Z. Liu et al. 2020; Lees et al. 2020; Kavvas et al. 2018; Sunuwar and Azad 2021; Mieth et al. 2016; Noman et al. 2023). These approaches vary in terms of intended scope and WGS data preprocessing. More complex, nonlinear, deep learning based approaches have also been developed for predicting AMR (Arango-Argoty et al. 2018; Mieth et al. 2021). While deep learning based approaches have more discriminative power than linear models, their predictions are less easily interpreted by humans (Guidotti et al. 2018). In many contexts, the interpretability of these AMR prediction models is a desirable feature, as it allows users to identify the genomic loci most responsible for positive and negative predictions. These identified loci can then be further studied to gain a deeper understanding of the predicted associations.

All such methods require genomic sequence data, which may not be available or practical in some circumstances, and predictions made from such models will include more uncertainty than direct phenotypic testing. In this sense, laboratory AST data is more robust for direct application in clinical settings. As AST is a cornerstone of routine clinical microbiology and AMR surveillance, it is simpler to acquire than physical isolates and subsequent WGS. AST data from surveillance are often available in a magnitude beyond that which can be practically sequenced, allowing for the study of large populations. For this reason, we introduce a method which leverages ML-based techniques and existing genomic data to predict genomic information from larger, more widely available AST data. An existing longitudinal genomic collection of *E. coli* from BSIs (Gladstone et al. 2021) provided an opportunity to train a model to predict the MDR ST131 C clades from AST data and apply this to the freely available data from surveillance of AMR in UTIs in Norway. We set out to predict the burden of ST131-C in UTIs over the last decade from the *E. coli* UTI susceptibility data, to help understand the current trend of increasing ST131 and MDR in BSIs.

## 6. Methods

### 6.1 Study Design and Data

This study utilised a longitudinal genomic dataset of 3254 *E. coli* BSI isolates with associated ST and AST data (Gladstone et al. 2021). These isolates represent an unbiased selection of approximately 15% of the larger Norwegian surveillance program on resistant microbes (NORM) between 2002-2017. NORM had collected all BSI isolates within 6 months, and all UTI isolates within 2-5 days each year between 2007-2021. 50 BSI and UTI isolates were collected per year <2007. (“NORM NORM-VET Reports” 2024) AST measurements were available for 24866 *E. coli* BSI and 28639 *E. coli* UTI isolates spanning 2006-2021. The AST test results for each isolate include disk diffusion zone diameters for multiple antimicrobials. Two different methodologies generated the AST data; from 2006-2010 by semi-confluent growth and from 2011 onwards according to the European Committee on Antimicrobial Susceptibility Testing (EUCAST) disc diffusion method (ESCMID-European Society of Clinical Microbiology and Diseases 2025; Matuschek, Brown, and Kahlmeter 2014).

The antimicrobials of interest were reduced to ciprofloxacin, gentamicin, and ceftazidime. These were chosen due to their relevance for the MDR ST131 C clades (C1 and C2), the availability of AST results in both the BSI and UTI AST data, and because they are non-overlapping with respect to antibiotic classes.

All datasets were divided into samples from before 2011 and from 2011 onwards. This was done to separate the data before and after the change in methodology. The number of available data points in each group is summarised in Table 1.

**Table 1.** Summary of dataset sizes for the sequenced BSI and UTI isolates as well as the unsequenced BSI and UTI datasets.

| Dataset | 2006-2021 | pre-2011 | post-2011 |
| --- | --- | --- | --- |
| BSI (genomic + AST) | 2790 | 813 | 1977 |
| BSI (AST only) | 24866 | 5441 | 19425 |
| UTI (genomic + AST) | 0 | 0 | 157 |
| UTI (AST) | 22942 | 7466 | 15476 |

For the genomic BSI dataset, ST was determined by using SRST2 v.0.2.0 (Inouye et al. 2014). ST131 clades were assigned based on clade-specific SNPs and *fimH* alleles (Ben Zakour et al. 2016; Price et al. 2013; Roer et al. 2017), and corrected based on the phylogenetic structure within the popPUNK lineage corresponding to ST131 published previously (Lees et al. 2019; Gladstone et al. 2021).

The AST data for 157 Norwegian *E. coli* UTIs isolated in 2019 from hospitalised (n=116) and community (n=41) patients at Akershus University Hospital were shared by Handal, Kaspersen *et al.,* paired to their published genomic dataset (Handal et al. 2025), which was used for further validation. The ST and clade membership of these isolates were determined using the same methods described above for the genomic BSI data. The AST distributions seen in this UTI sample set were compared to the BSI AST distributions from 2013 onwards using a multivariate Cramér test to determine if the underlying distributions were significantly different. This test was implemented in the cramer package v0.9-4 in R. The Cramér test was also used to compare ST131-C isolates within the BSI dataset to all others in order to determine if there is a statistically significant difference in the multivariate AST distribution. This test was performed using the genomic BSI dataset, and three tests were performed: one comparing ST131-C to all other isolates across all years, once for isolates from before 2011, and a third for isolates from 2011 and onwards.

### 6.2 Classifier Training

Classifiers were trained for predicting isolate membership to the ST131-C within the *E. coli* population. Logistic regression, Random Forest, and XGBoost classifiers were implemented in the python packages scikit-learn v1.5.1 and xgboost v2.0.3. Random Forest and XGBoost model types were chosen due to their relative simplicity and speed of training, while still being capable of learning nonlinear decision boundaries. The logistic regression model is included as a comparative baseline for a simple linear classifier. All models were defined as binary classification models predicting inclusion or exclusion from the pre-defined population group, ST131 clade C. This inference is made using the disk diffusion zone diameter values for ciprofloxacin, gentamicin, and ceftazidime measured in millimetres.

The genomic BSI dataset was split into a training and test set by selecting 75% of the dataset for training (n=2093) and setting aside the remaining 25% for testing (n=698). This split was stratified by both isolation year and ST131-C membership to ensure both training and test sets had similar distributions for isolation year and training labels. To accomplish this stratified split, only one ST131-C isolate from 2006 in the genomic BSI dataset was discarded. The training data was used to train the two models using a stratified 5-fold cross-validation. The Random Forest and XGBoost classifier hyperparameters were chosen by selecting the best-performing model over a grid search. This performance was evaluated by selecting the highest mean F1-score across the 5 cross-validation folds. The test set was used to estimate classifier performance metrics, including classification accuracy, sensitivity, and specificity.

As methods for disk-diffusion-based AST measurements were altered in 2011, the dataset was also partitioned into samples from before and after 2011. These two data groups were also independently divided into training and test sets following the 75/25 split described above. The same training methods described above were used to train models on the pre-2011 and 2011-onwards data independently from one another. Consequently, for each model type, 3 unique instances were trained: one with data from all years, a second with data from before 2011, and a third with data from 2011 and onwards. The latter two were used in combination for subsequent prediction and evaluation, only predicting isolates originating from the years they were trained on, which we will refer to as the split-year method.

### 6.3 Prediction

The classifiers trained using the split-year BSI genomic plus AST dataset were utilised with the disk diffusion zone diameter values for isolates in the UTI dataset of 22942 and BSI dataset of 24866 unsequenced *E. coli* isolates to predict their membership in ST131-C. The predictions for isolates taken within each year were used to estimate the total fraction of recorded samples that belong to ST131-C.

False discovery and false omission rates were estimated using the holdout test samples and then used to compute error bounds for each year’s prediction. The expected false discovery rate (FDR) was used to approximate the number of positive predictions that are type I errors. This approximation was used to set a lower bound on the number of positive predictions for the group of samples within a year. The same procedure using the false omission rate (FOR) and type II errors was implemented to derive an upper error bound for the positive predictions. These values are determined using the equations shown below.

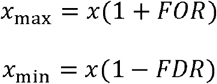

The yearly prevalence trends in the test dataset were compared to the ground truth trends using a Pearson correlation coefficient. This correlation statistic was used to conduct a hypothesis test for a positive correlation between the true and predicted trends. A permutation test with 50000 independent resamples was used to compute a p-value.

## 7. Results

### 7.1 Trained Classifiers act as predictors for ST131-c membership

The models trained on each of the 5 training cross folds were evaluated on the withheld BSI test data samples (n=698) with both known ST131-C membership and AST data. The computed metrics are shown in Table 2. From the test performance, the split-year trained models achieved the best F1-scores, specificity, and precision. ST131-C is relatively uncommon in the test set at only 30 of the 698 samples. This large data imbalance makes accuracy and specificity incomplete performance metrics, so we focus more on the F1-scores and precision values of each model. The classifier prediction performance aligns with Cramér tests comparing ST131-C AST distributions to the rest of the population. Tests utilizing the entire dataset, only isolates from before 2011, and only isolates from 2011 onwards all found ST131-C AST distributions to be significantly different from the rest of the *E. coli* BSI population with estimated p-values indistinguishable from 0.

**Table 2.** Mean and standard deviation of performance metrics of models trained on 5 training cross folds. The metrics shown were computed by evaluation against test samples withheld from all training folds.

| Model | Training Method | F1-Score | Specificity | Sensitivity / Recall | Precision |
| --- | --- | --- | --- | --- | --- |
| Linear | Combined | 0.606 ± 0.000 | 0.942 ± 0.000 | **1.000 ± 0.000** | 0.435 ± 0.000 |
|  | Split | 0.636 ± 0.006 | 0.948 ± 0.002 | 0.994 ± 0.014 | 0.468 ± 0.008 |
| Random Forest | Combined | 0.615 ± 0.061 | 0.980 ± 0.003 | 0.647 ± 0.077 | 0.588 ± 0.054 |
|  | Split | 0.658 ± 0.027 | **0.981 ± 0.002** | 0.697 ± 0.037 | **0.625 ± 0.028** |
| XGBoost | Combined | 0.649 ± 0.017 | 0.972 ± 0.004 | 0.780 ± 0.030 | 0.556 ± 0.028 |
|  | Split | **0.687 ± 0.015** | 0.975 ± 0.004 | 0.807 ± 0.032 | 0.600 ± 0.030 |

The false positives in the test set for all classifiers listed above were fairly evenly distributed among different STs. Three of the 15 BSI false positives from the split-year trained XGBoost model predictions belonged to ST405, two from ST90, and the rest belonged to unique STs. Predictions from the split-year trained random forest model displayed 2 of the 13 false positives from ST90 and ST405, with the rest being from separate STs. For the six false negative CC131-C isolates, there was no phylogenetic clustering; they were spread sporadically across the phylogeny. Additionally, the distributions of the most common population groups’ zone diameter measurements were visualized to verify that none of the major STs overlapped significantly with ST131-C (Figure 1). ST131-C has a very distinct distribution across the three antimicrobials, especially in ciprofloxacin resistance. The only ST that overlaps at all with the ciprofloxacin disk diameter values of ST131-C is ST393, which differs enough in its gentamicin and ceftazidime disk diameter distributions such that it is not a major confounding ST for the predictive models. The zone diameter distributions in the most common STs within the genomic UTI dataset are also visualized (Supplementary Figure S1) and show no specific STs that overlap entirely with ST131-C. Within these sequenced UTI isolates, the ciprofloxacin and gentamicin distributions of ST131-C are very unique, while ST38 and ST384 show some overlap with ST131-C in ceftazidime zone diameter values.

**Figure 1.**
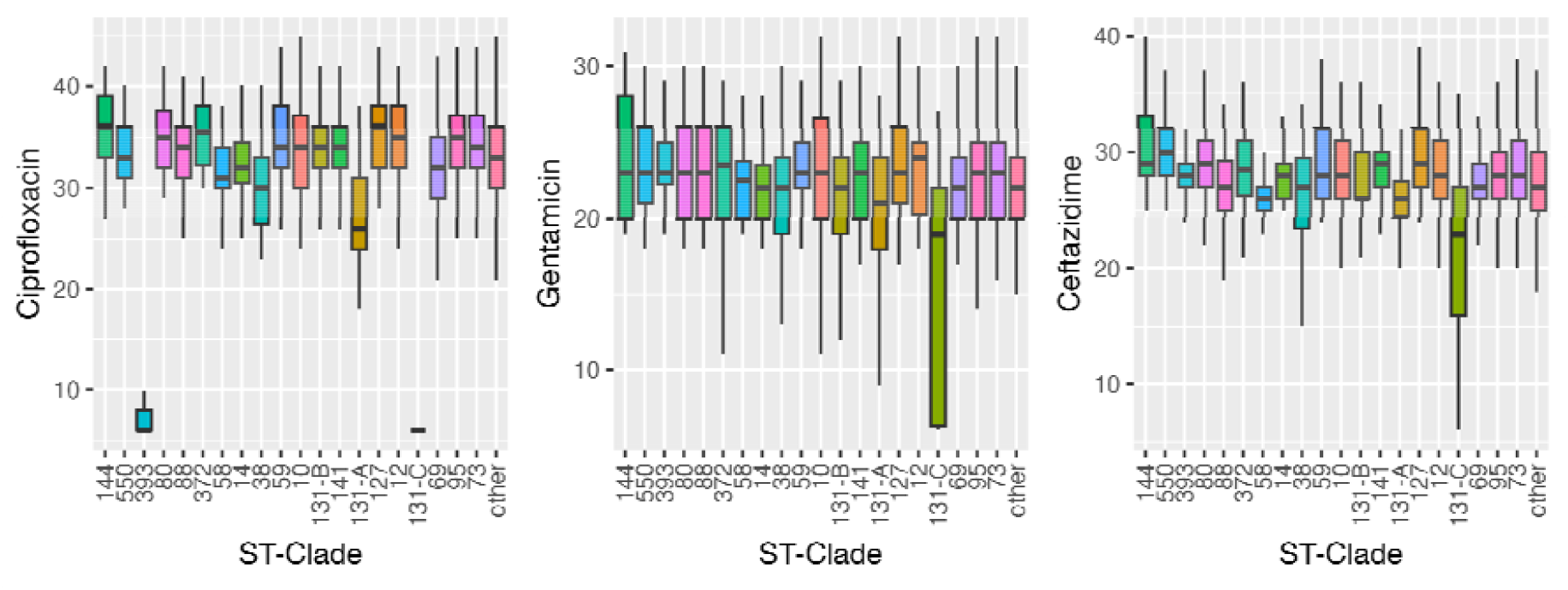
Boxplots for the ciprofloxacin, gentamicin, and ceftazidime zone diameter distributions in the top 20 most common population groups in the sequenced NORM dataset.

The split-year models were further validated using the 157 UTI AST profiles shared by Handal, Kaspersen et al., which were paired with the genomic data (Handal et al., 2025). Within this group, 7 isolates belonged to ST131-C. These 7 isolates were compared to the ST131-C BSI isolates and could not reject the null hypothesis that the UTI and BSI ST131-C isolates follow the same distribution (p=9.690). This is further discussed in the supplementary section S2. The mean value of ciprofloxacin, gentamicin, and ceftazidime zone diameters of these 7 isolates were 6, 12, and 20 mm, respectively. These are similar to the BSI means of 6, 15, and 21 mm. We did not perform a hypothesis test to compare these distributions due to the low number of sequenced ST131-C isolates from UTIs. The baseline linear classifier correctly classified all 7 of these isolates as ST131-C, as well as 8 other isolates (all from unique STs). The random forest model correctly classified 4 ST131-C isolates and predicted no false positives, the rest of the ST131-C were incorrectly predicted as other isolates. The XGBoost model correctly predicted all 7 ST131-C isolates, along with 1 false positive from ST131 clade A.

The BSI training and test data samples were also used to evaluate the reliability of the trained models for estimating the yearly prevalence of ST131-C. These estimates, along with the ground-truth values for the sequenced data, are shown in Figure 2.

**Figure 2.**
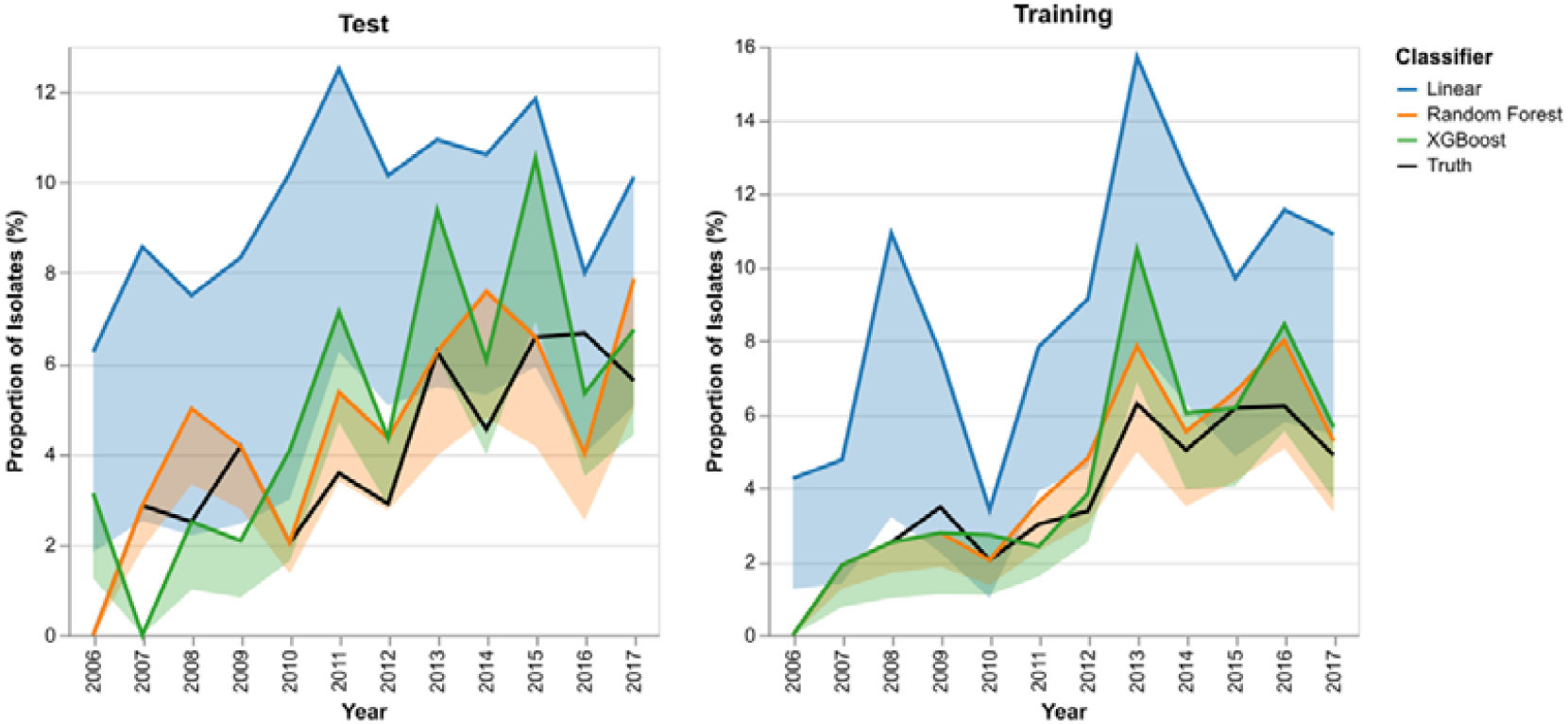
Yearly predicted and ground-truth prevalence of ST131-C in the sequenced dataset. Predicted trends derived from split-year trained random forest and XGBoost models.

To more thoroughly investigate the quality of the fits in Figure 2, the Pearson correlation coefficient was calculated between each of the predicted trends and the ground-truth fractions. The calibration plots are shown in Figure 3.

**Figure 3.**
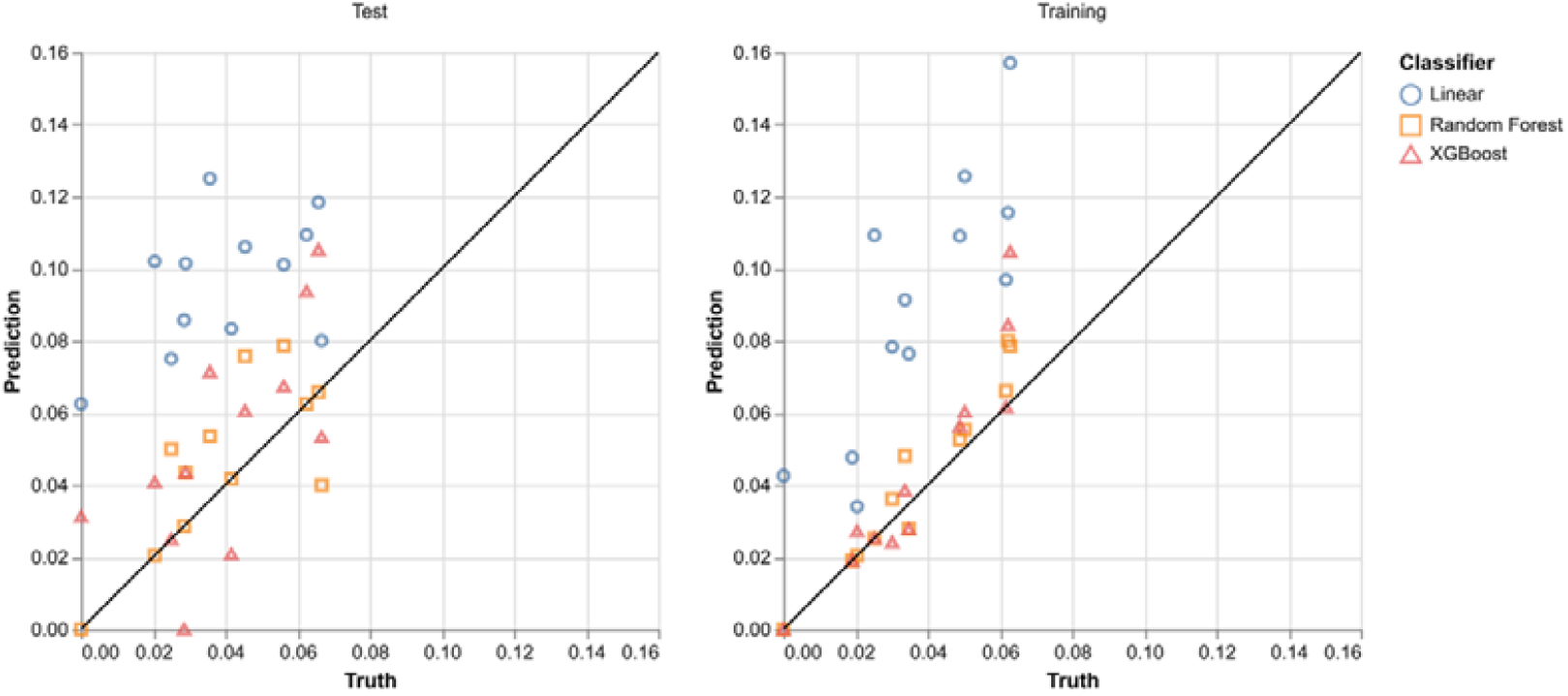
Calibration plot between predicted yearly fraction values and the true yearly fractions in the test and training datasets. Predicted values are derived from split-year trained random forest and XGBoost models.

The Pearson correlation (*r*) coefficient for the test predictions using the Random Forest and the XGBoost classifiers are 0.7461 and 0.6716, respectively. Hypothesis tests using a null hypothesis of no correlation (*r*=0) and an alternative hypothesis of positive correlation (*r*>0) were computed. The resulting p-values were 0.0036 and 0.0076 for both the Random Forest and XGBoost models, respectively. These indicate a positive correlation between the true and predicted yearly prevalence values of ST131-C E. coli in Norwegian BSI data.

The qualitative fit of the model predictions on test data in Figure 2, as well as the positive correlations between truth and predicted yearly prevalence, are evidence of little overfitting.

### 7.2 Random Forest feature importance values reflect sensitivity to testing procedures

Random Forest models offer useful insight for analyzing the importance of each feature used for prediction. For all data ranges, the importance of the antibiotics in the final model aligns with the corresponding mean and standard deviation of the corresponding cross-folds. Notably, the antibiotic importance values of the models trained on data from 2006-2010 don’t align with the other two data ranges (Table 3). This is likely a result of the distribution shift caused by the change in laboratory testing procedures. This distribution shift is more explicitly visible in Figure 4, where the modes of gentamicin and ceftazidime zone diameter distributions decrease between 2010 and 2011. The median gentamicin zone diameter drops from 26mm to 22mm between 2010 and 2011, ceftazidime drops from 32mm to 27mm, and ciprofloxacin drops from 37mm to 34mm.

**Table 3.**
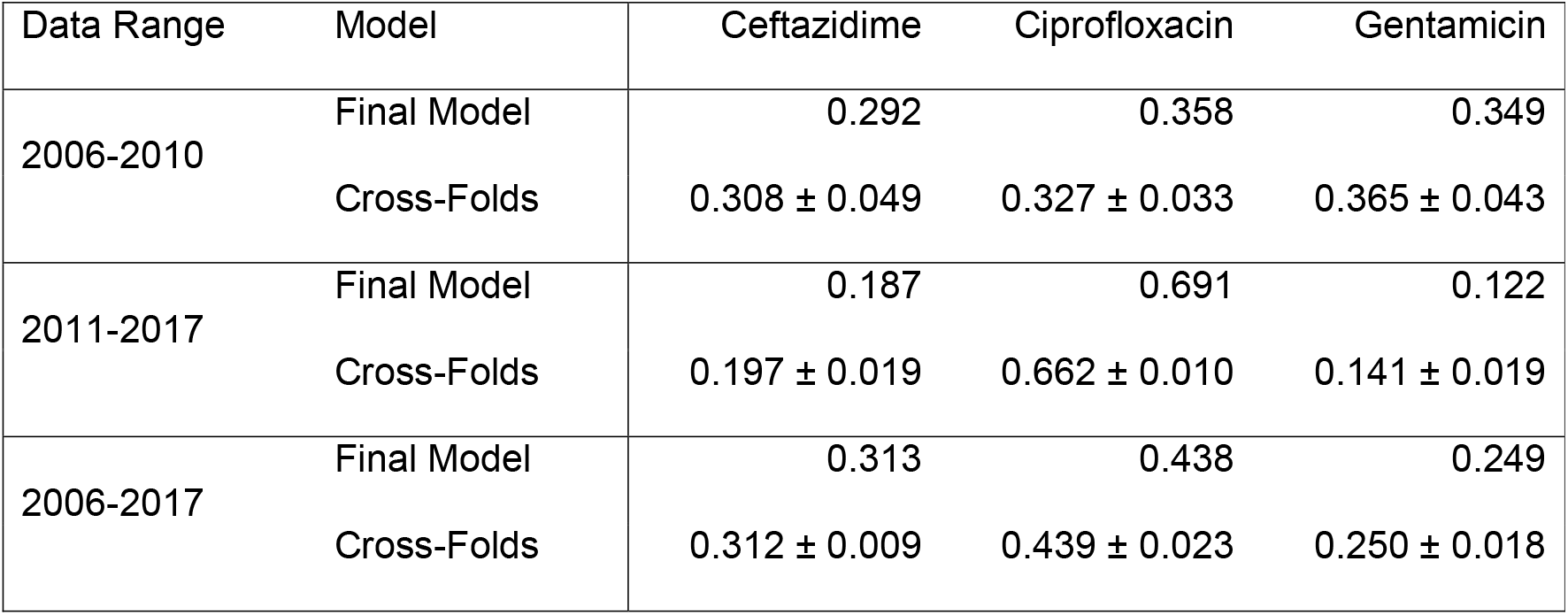
Importance of each antibiotic in the trained random forest models.

| Data Range | Model | Ceftazidime | Ciprofloxacin | Gentamicin |
| --- | --- | --- | --- | --- |
| 2006-2010 | Final Model | 0.292 | 0.358 | 0.349 |
|  | Cross-Folds | 0.308 ± 0.049 | 0.327 ± 0.033 | 0.365 ± 0.043 |
| 2011-2017 | Final Model | 0.187 | 0.691 | 0.122 |
|  | Cross-Folds | 0.197 ± 0.019 | 0.662 ± 0.010 | 0.141 ± 0.019 |
| 2006-2017 | Final Model | 0.313 | 0.438 | 0.249 |
|  | Cross-Folds | 0.312 ± 0.009 | 0.439 ± 0.023 | 0.250 ± 0.018 |

**Figure 4.**
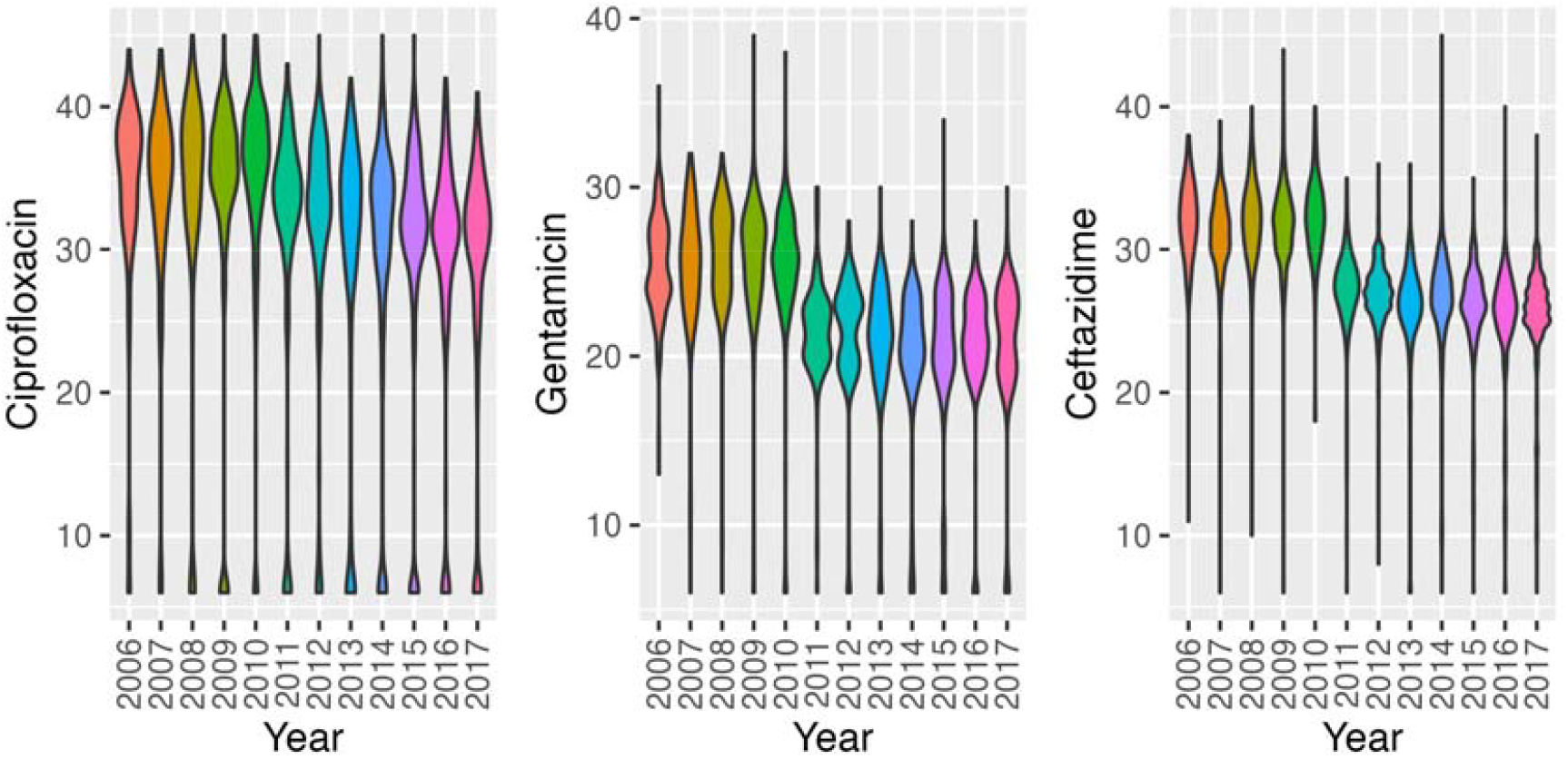
Violin plots of the distribution of disk diffusion zone diameters for each year within the sequenced BSI population.

### 7.3 Model predictions show a steep increase in ST131-C in both BSI and UTI isolates

The trained XGBoost split-year models were used to predict yearly prevalence of ST131-C in the unsequenced UTI and BSI isolates. Figure 5 displays the trend of this prevalence as predicted by split-year models. The estimated trend shows a sharp increase in ST131-C prevalence in 2011.

**Figure 5.**
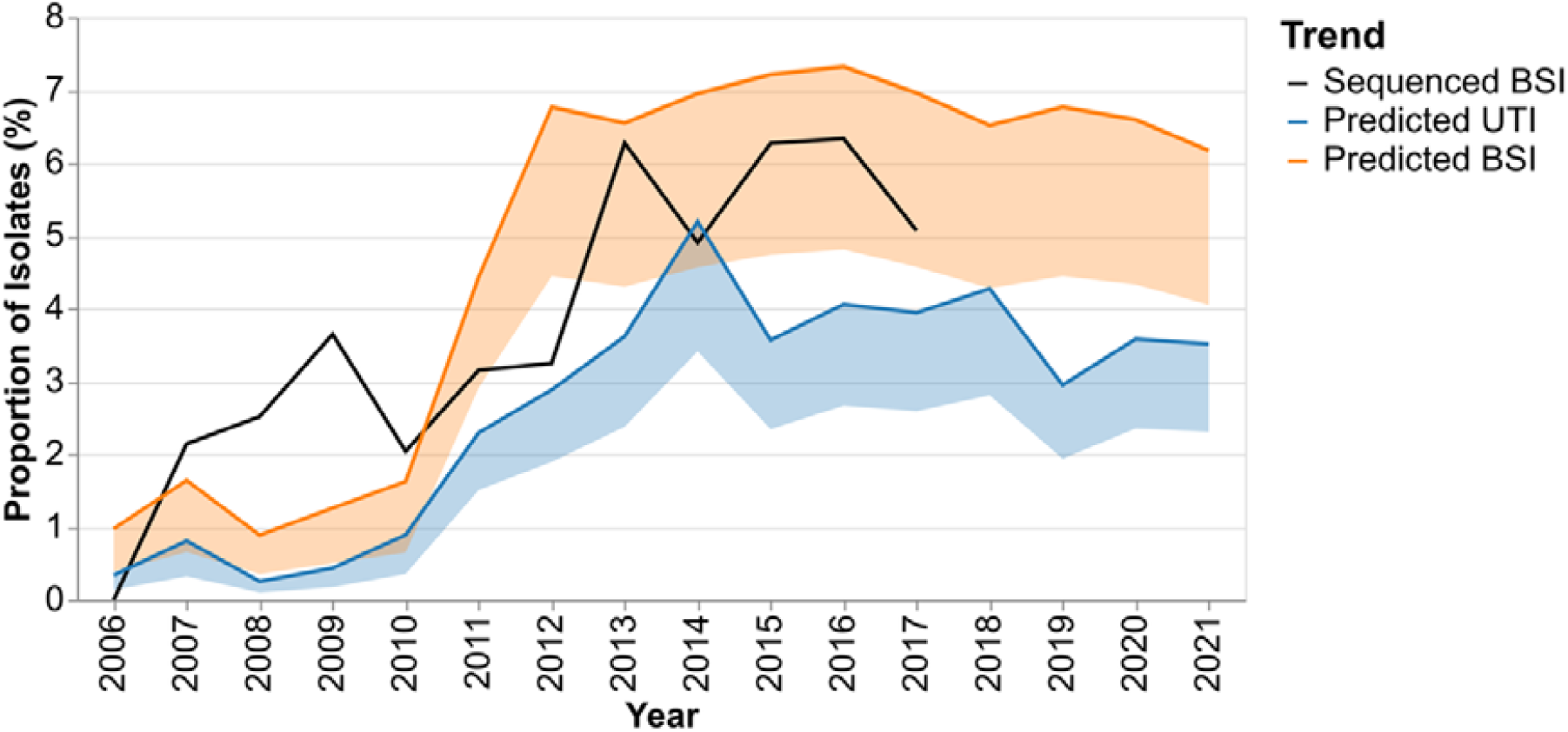
Predicted yearly prevalence of ST131-C in the unsequenced UTI and BSI AST datasets using the XGBoost classifier models trained on pre- and post-2011 data separately. The trend from the sequenced BSI data is plotted for comparison.

The BSI genomic data shows peaks relative to years before and after 2009, 2013, 2015, and 2016, but the genomic dataset was 1/10 of the size of the BSI AST dataset, and the absolute number of ST131-C was small (n=121), which can introduce some noise to the yearly prevalence, especially when the prevalence was lowest before 2013. Predictions show a lower prevalence of ST131-C in UTIs than in BSIs. This is reflected by multivariate distributions of the zone diameter measurements for UTI and BSI populations being significantly different (p=9.99×10-4). Across all years shown, the mean ratio of UTI to BSI ST131-C prevalence is 0.5054 (SD 0.1176). All three datasets show an increase in ST131-C prevalence between 2010 and 2014, followed by a stable prevalence in the AST datasets. Data on the relative prevalence of ST131-C for the period 2014-2017 in carriage (1.7%) and disease (12.6%) from the UK (Gladstone et al. 2024; Pöntinen et al. 2024) allowed us to infer the likely prevalence of carriage in Norway in this same time period would be 7.4 times lower than BSIs and determine the relative ST131-C enrichment between all three niches (carriage, UTI, BSI). An inferred ST131-C carriage prevalence of 0.95% in Norway was 4.0 times lower than in UTIs (4.1% AST XGBoost predictions), whilst the UTI prevalence was 1.7 times lower than in BSIs (7.1% AST XGBoost predictions, 7.0%, genomic collection) between 2014 and 2017. This indicates that a greater enrichment of ST131-C between carriage and BSI occurs in the progression from colonisation to UTIs than from UTIs to BSIs. However, both represent disease progression where ST131-C increases more than its competitors.

## 8. Discussion

The XGBoost model displayed the most reliable test performance, displaying the best F1-score and sensitivity values on the unbalanced test dataset. All the trained predictive models exhibited very high accuracy and specificity values of over 96 percent, which is likely due to this imbalance. The specificity and sensitivity values suggest the models can offer upper estimates of ST131-C membership. This manifests as a higher probability of producing type I errors (false positive) than type II (false negative). Critically, the specificity, sensitivity, and correlation scores of the yearly trends indicate that they are more robust for predicting population-wide trends in ST131 clade C prevalence, which we present here, rather than predicting individual isolates.

The only misclassification from the XGBoost model in the genomic UTI dataset belongs to ST131 clade A, displaying notably better performance than observed in the genomic BSI dataset, as shown in Table 2, which supports the observation that this UTI dataset does not contain a specific population group with AST distributions overlapping with those of ST131-C. Additionally, the AST distributions of ST131-C in both UTI and BSI overlap enough to reliably use the BSI-trained models to predict UTI isolates. ST38 overlapped with the ST131-C ceftazidime distributions in both UTIs and BSIs, and ST393 overlapped with the ciprofloxacin values of ST131-C in BSIs. There were no STs observed in either BSIs or UTIs that overlapped significantly with ST131-C in all three of the antimicrobials used.

The difference in performance between the BSI and UTI datasets may be driven by the reduced temporal and spatial diversity in the genomic UTI data. The genomic UTI isolates used for this verification were isolated from a single laboratory(Akershus University Hospital) in 2019, while the sequenced BSI isolates originate from 15 laboratories across Norway between 2006 and 2017. Another possible explanation for the disparity in classifier performance between UTI and BSI isolates is the potential diversity of MDR clones causing each infection type. If the BSI causing *E. coli* within Norway includes a wider array of MDR clones beyond ST131-C, they would contribute to higher false positive rates within the BSI population. However, the previously mentioned difference in temporal and spatial diversity could produce the difference in MDR clone diversity. More thorough genomic studies of the relationships between UTI- and BSI-causing *E. coli* would be required to further understand this.

In the genomic BSI dataset, the false positives are dispersed among several different clones. The genomic UTI dataset used for model validation contained too few false positives from the XGBoost and random forest models to understand the ST distribution of potential false positives. The false positives in the genomic UTI dataset from the linear model were also evenly distributed among multiple STs. These results suggest that there are no specific confounding STs within our training and test datasets. However, it is still possible for an MDR clone with a similar AST profile to ST131-C to increase false positives when making inferences with unsequenced isolates.

The antibiotic classes and mechanisms that underlie resistance for the three antibiotics used to predict ST131-C differ. Ciprofloxacin resistance is predominantly chromosomally mediated. In contrast, the genes contributing to gentamicin and ceftazidime resistance are typically located in mobile genetic elements (MGEs), which can simultaneously carry numerous antimicrobial resistance genes (ARGs) (Park 2014; Miró et al. 2013). These mobile ARGs can be transferred into other clones; however, the exact level of resistance, as measured here by the diffusion zone, also differs by the lineage in which the gene is expressed (Card et al. 2021). By including an antibiotic with chromosomally encoded resistance, we reduce the likelihood that the same profile is found in other clones as the result of a shared MGE. The ST131-C ciprofloxacin disk diffusion zone of 6 mm was also rare in the BSI collection (6.8%) and particularly indicative of ST131-C (51%, Figure 1). The prediction of ST131-C depended on its dominating a relatively rare AMR profile and the granularity provided by the disk diffusion zone; such predictions would not be possible for more common combinations of resistance. Whilst some false positives reduce the specificity for ST131-C, they nonetheless represent an underlying AMR profile of clinical concern and contribute to a worrying trend.

The UTI ST131-C prediction results show a significant spike in prevalence in Norway from 2011. The growth of ST131-C appears to plateau after a few years for UTI and BSI predictions and is stable for almost a decade, remaining between 3 and 4 percent of all UTIs (Figure 5). The sequenced UTI data from 2019 is slightly higher, with ST131-C prevalence in that year at 4.3%, though it represents only a single region near the capital of Norway. This suggests some form of equilibrium point had been reached. These observations align with growth plateaus observed in the estimated population size for ST131 clades (Gladstone et al. 2021) and the findings of an epidemiological study modeling the infection dynamics for the NORM *E. coli* BSI data used in this study. It was shown that clade C2 exhibits a significantly lower R0 than clade A isolates (Ojala et al. 2024), suggesting it isn’t as good at colonising or spreading. Despite being poorer spreaders, clades C2 and C1 are more likely to cause disease; they have higher frequencies in BSIs than expected, given the exposure frequency in healthy colonisation (Gladstone et al. 2024; Pöntinen et al. 2024; Mäklin et al. 2022). Here, we observe that ST131-C prevalence in UTIs is lower than in BSIs across all years. This suggests that the overrepresentation of ST131-C in BSIs is not solely due to its propensity to cause UTIs with a proportional overspill into BSIs. This relationship is consistent with the respective prevalence of ESBL-positive isolates in UTIs and BSIs in the NORM reports for the sample time period (“NORM NORM-VET Reports” 2024). However, our model has approximately half as many false positives and negatives as using ESBL status alone. Hard-to-treat UTIs will have a greater opportunity to cause a BSI, and ST131-C *E. coli* may possess further virulence factors that facilitate invasion and survival in the human bloodstream compared to other UTI clones. It must be noted that whilst the urinary tract is the dominant focus for BSIs in Norway (67% (Mehl et al. 2017)), other routes of infection (e.g biliary tract) will also influence the trends in BSI.

The analysis of Norwegian UTI populations presented are conducted using AST data exclusively to infer genotypic information on the clonal membership of these *E. coli* infections. This method allowed for a broad-scale analysis and prediction of over 40000 bacterial isolates without requiring dedicated genotypic testing. While the accuracy of an ML prediction-based epidemiological study will be limited compared to one performed with molecular methods such as WGS or sequence typing or PCR, the accessibility of AST data allows for the application of retrospective studies at large scale with very low costs. The predictions also allow for targeted sequencing for in-depth follow-up, a resource-efficient alternative to population genomics.

The models used in this study are very specifically trained on a limited scope of potential AST data. ST131-C is a subpopulation of *E. coli* with characteristic and problematic MDR patterns. Consequently, the models trained and used in this study are highly specific to predicting membership in this particular subpopulation from *E. coli* disk diffusion measurements. Because of this, the shared code is an example of how such a study can be performed and not a generalizable tool to be applied to other study conditions.

The predictive power in differentiating population groups would increase with the inclusion of more relevant and informative antimicrobials where available. The models implemented in this study utilised only 3 informative antimicrobials from different classes. Additional antimicrobial substances may provide additional discriminative power, but only if they are relevant to the specific task and microbe at hand and do not overlap entirely with the resistance profiles of the already included antimicrobials. For example, the inclusion of another fluoroquinolone with chromosome-associated resistance factors might only add redundant information to the AST profile. As such, the antimicrobials used for these predictive models require careful consideration and must be selected leveraging existing biological and clinical knowledge.

The type of AST data used also has a consequence for model training. This study used zone diameters measured in millimetres of disk diffusion tests. This specific measurement has the advantage of being recorded on a linear scale, which improves training stability depending on the model chosen. In contrast, MIC tests increase exponentially as powers of 2. These MIC values, when available, are still highly valuable data to collect and would likely still be useful for training models such as following this study’s procedure. A log_2_ transformation applied to MIC measurement data would maintain monotonic relationships while creating a more linear scale for the training. We observed a shift in the available data and learned model weights corresponding to a standardised change of laboratory testing practices established in 2011. The distinct shift in the weights attributed to each antibiotic indicates that laboratory practices heavily influence data and, therefore model training.

Of note, due to biological relatedness of data samples, applications of predictive ML algorithms in biological settings are prone to data leakage if it is not explicitly controlled for (Bernett et al. 2024). This can often result in test results overestimating a model’s performance and generalizability. A common way to control for this is to divide datasets according to population cluster boundaries or by phylogenetic groups, intentionally excluding multiple clusters from training. These methods better emulate how the model may perform when encountering completely novel samples. However, the models trained in this study are explicitly attempting to predict a sample’s membership to a population group, meaning this particular data leakage concern is not relevant to the practices used here.

The AST data used in this study is more accessible and widely available than the genomic sequence data used for population clustering and clade identification, as shown by the disparity in sizes of the genomic BSI and the AST BSI and UTI datasets in this study. Laboratory susceptibility tests are often performed routinely at scale in many countries globally. The method proposed in this study could allow for large-scale epidemiological studies to be performed in settings without continuous access to sequencing technology, the ability to store isolates, or with historical data that is no longer possible to sequence. This would open the possibility of improved public health studies in economically underprivileged regions. However, this would still require some amount of genomic sequencing to be done in the said resource-limited settings in order to acquire relevant and informative training data, as it should not be expected that the models we trained on Norwegian *E. coli* isolates would generalise well to nations in Sub-Saharan Africa or Southeastern Asia, for example.

After an initial rapid expansion in ST131-C prevalence, its subsequent stable persistence between 2015 and 2021 in Norwegian UTIs and BSIs suggests that the clade has remained a stable competitor in its existing niche. Also, the endemic nature of its persistence further suggests that the inter-clone competitiveness of ST131-C in the urinary tract against other clones and clades of *E. coli* remains unaltered since its emergence. Yet, the enrichment of ST131-C in BSIs compared to UTIs suggests this clone can more easily progress to an invasive infection when compared to the rest of the population as a collective. It would thus be attractive to conduct similar studies for other regions. Understanding geographical differences in the bacterial population structure, the rate at which competitive MDR lineages can enter a niche, their subsequent endemic prevalence and disease progression rates could open the door for vaccine-based manipulation of the population structure of commensal *E. coli* to prevent disease progression from the gut.

## 9. Author statements

### 9.1 Author contributions

T.A.R wrote the shared code, trained the machine learning models, performed predictive analysis with said models, and primarily wrote the manuscript alongside R.A.G.

A.K.P., E.H., M.K., and K.H. supervised the project and assisted with study design and manuscript editing.

Ø.S. provided resources and guidance on clinically relevant aspects, manuscript editing, and provided additional guidance.

J.C. assisted with study design, manuscript editing, and provided additional guidance.

R.A.G. conceptualised the study, provided guidance on biological and epidemiological aspects, and primarily wrote the manuscript alongside T.A.R.

All authors reviewed the manuscript.

### 9.4 Ethical approval

All data used in the study were anonymized. Thus, no ethical approval was required.

### 9.5 Consent for publication

No sensitive information is disclosed in this manuscript.

## 9.6 Acknowledgements

We are grateful to the Norwegian Surveillance System for Resistant Microbes (NORM) for making data available and to all Norwegian clinical microbiology laboratories for contributing data to NORM. We also thank Dr Handal, Dr Jørgensen and Dr Sunde for sharing the relevant accompanying antimicrobial susceptibility data for their Norwegian UTI genomic dataset.

