## Supplementary Material for "Machine learning based lineage prediction from AMR phenotypes for Escherichia coli ST131 clade C surveillance across infection types"

### S1. Supplementary Figures

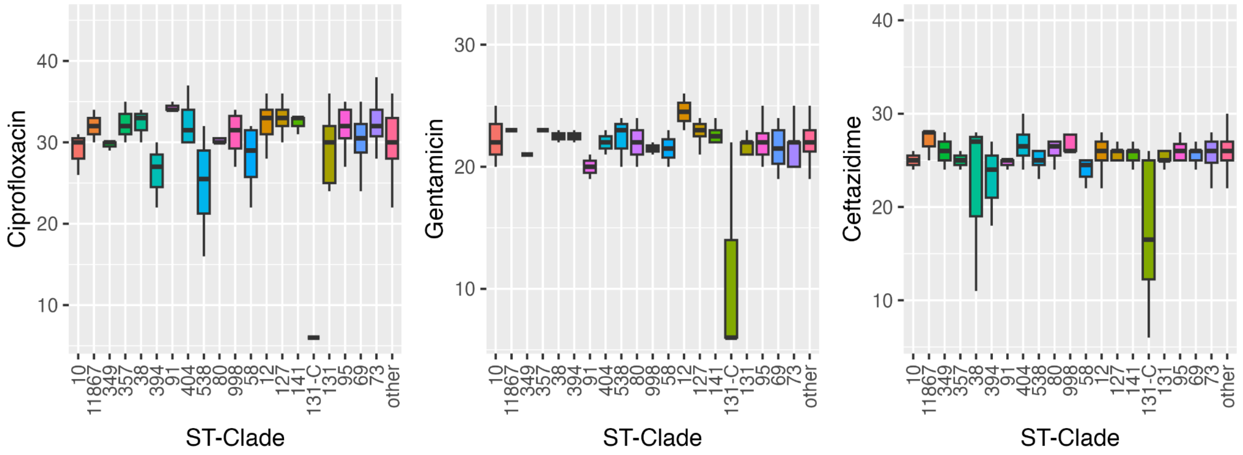


**Figure S1:** Boxplots for the ciprofloxacin, gentamicin, and ceftazidime zone diameter distributions in the top 20 most common population groups in the sequenced UTI dataset.

### S2. Comparing ST131-C isolates in BSIs and UTIs

While a Cramer test shows that ST131-C AST distributions between BSIs and UTIs are the same, this test was conducted with a small amount of ST131-C UTI isolates. This section explores the comparison between these groups more.

**Table S1:** Percentage of UTI isolates in respective subgroups fall between the first and third quartile of the zone diameter distributions of the ST131-C in the genomic BSI dataset. Here, the quartiles of each zone diameter distribution were computed independently of the other two antimicrobials.

| Antimicrobial | ST131-C | non ST131-C | ST394 | ST58 |
|---|---|---|---|---|
| Ciprofloxacin | 100.00 | 2.67 | 0.90 | 0.90 |
| Ceftazidime | 42.86 | 83.11 | 64.95 | 48.48 |
| Gentamicin | 28.57 | 56.00 | 29.21 | 17.11 |

Observing the zone diameter distributions individually shows significant overlap of non-ST131-C UTI isolates with the zone diameter values of ST131-C found in BSIs for Ceftazidime and Gentamicin but not Ciprofloxacin (Table S1). Every ST131-C UTI fell within the first and third quartiles of at least one of the ST131-C zone diameter distributions in the BSI population (Figure S2-B). Conversely, no non-ST131 isolates from the UTI population fell within these ranges for more than two of the three antimicrobials (Figure S2-A). This lack of complete overlap in the zone diameter distributions means each of these isolates can be distinguished from ST131-C in at least one of the three antimicrobial zone diameters. This also explains why the linear logistic regression classifier failed in more cases than the nonlinear classifiers.

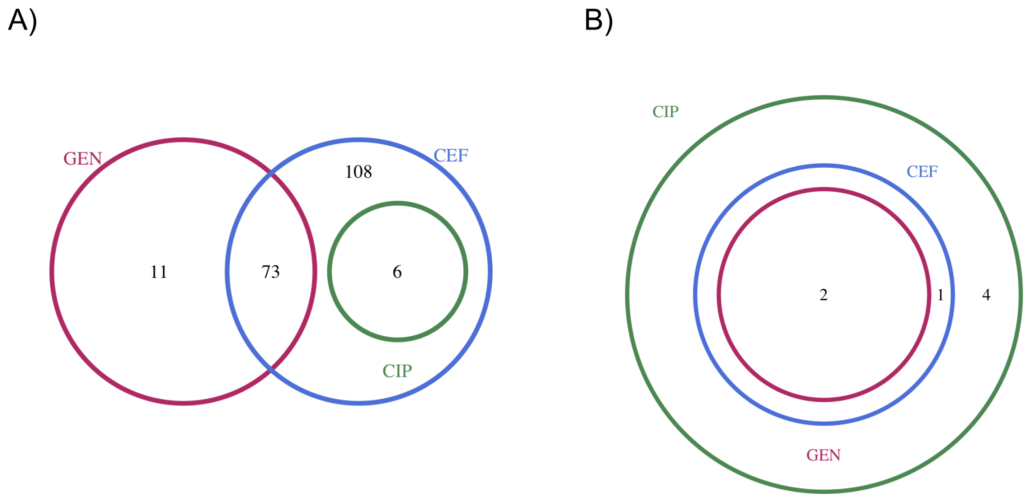

**Figure S2:** Venn-diagrams visualizing the isolates that fall between the first and third quartiles of the zone diameter distributions in the genomic BSI dataset from the  A) non-ST131-C UTI isolates that and B) ST131-C UTI isolates.
